# Target-template relationships in protein structure prediction and their effect on the accuracy of thermostability calculations

**DOI:** 10.1101/2022.09.19.508270

**Authors:** Muyun Lihan, Dmitry Lupyan, Daniel Oehme

## Abstract

Improving protein thermostability has been a labor- and time-consuming process in industrial applications of protein engineering. Advances in computational approaches have facilitated the development of more efficient strategies to allow the prioritization of stabilizing mutants. Among these is FEP+, a free energy perturbation implementation that uses a thoroughly tested physics-based method to achieve unparalleled accuracy in predicting changes in protein thermostability. To gauge the applicability of FEP+ to situations where crystal structures are unavailable, here we have applied the FEP+ approach to homology models of 12 different proteins covering 316 mutations. By comparing predictions obtained with homology models to those obtained using crystal structures, we have identified that local rather than global sequence conservation between target and template sequence is a determining factor in the accuracy of predictions. By excluding mutation sites with low local sequence identity (<40%) to a template structure, we have obtained predictions with comparable performance to crystal structures (R^2^ of 0.67 and 0.63 and an RMSE of 1.20 and 1.16 kcal/mol for crystal structure and homology model predictions, respectively) for identifying stabilizing mutations when incorporating residue scanning into a cascade screening strategy. Additionally, we identify and discuss inherent limitations in sequence alignments and homology modeling protocols that translate into the poor FEP+ performance of a few select examples. Overall, our retrospective study provides detailed guidelines for the application of the FEP+ approach using homology models for protein thermostability predictions, which will greatly extend this approach to studies that were previously limited by structure availability.

## Introduction

Protein engineering for thermostability has a wide range of applications in chemical, pharmaceutical, and biotechnological industries^1,2^. In these industries, protein stability usually requires optimization to enable enzymes to withstand specific chemical conditions or to extend the shelf-life of protein products^3–5^. Directed evolution and phage display technologies have proven an effective optimization strategy for identifying stable variants^6–8^, however, can become laborious when multiple iterative rounds of optimization are needed. On the other hand, computationally-enabled rational design, based on methods that use protein sequence or structural information to computationally predict, rank, and thus screen for potential stabilizing mutations^9^, can greatly reduce the cost of identifying stabilizing mutants.

Various computational approaches for predicting protein thermostability have been developed over recent years using a combination of statistical, physical, and machine-learning methods^10,11^. As most experimentally validated mutations are destabilizing, one of the common major drawbacks of currently available machine-learning-based algorithms lies in the prediction bias caused by the use of skewed training datasets in model building^12–16^. In comparison, methods that use a physics-based description of proteins, such as Molecular Mechanics/Generalized Born Surface (MM/GBSA)^17^ and Free Energy Perturbation (FEP)^18,19^, can circumvent this reliance on fitting to such unbalanced experimental datasets. These methods rely on accurate force field descriptions of proteins and using a thermodynamic cycle involving both the folded and unfolded states of the wild-type and mutant forms of the protein to enable free energy estimates of changes in protein thermostability upon a mutation (ΔΔG)^20^. Combining these two methods in a cascade screening strategy has been proposed to reduce computational cost whilst also maintaining prediction accuracy^20^.

An implementation of FEP developed by Schrödinger, FEP+, has shown unparalleled accuracy in predicting protein thermostability upon single-point mutations in several retrospective studies. First applied to a dataset of 741 mutations, Steinbrecher et al., demonstrated an overall root mean square error (RMSE) of 2.27 kcal/mol and an R^2^ of 0.40^21^. At the same time, Ford and Babaoglu predicted protein thermostabilities with FEP+ on a dataset with melting temperature data from 62 mutations and produced an R^2^ of 0.50^22^. Owing to recent improvements in the force field^23^; water sampling around mutation sites^24^; protocols for simulating charge-changing mutations^25^, proline mutations^26^, and mutations that need additional consideration for different protonation states^27^, FEP+ achieved significant improvement in performance in two recent studies^20,28^. Duan et al. proposed a tripeptide model for better representation of the unfolded states of a protein and when applied to a dataset of 87 mutations, showed an improved RMSE of 1.11 kcal/mol and an R^2^ of 0.66^20^, on par with the accuracy achieved in protein-ligand binding free energy prediction^29^. Scarabelli et al. later extended the application to a larger dataset with 328 mutations and obtained an RMSE of 1.25 kcal/mol and an R^2^ of 0.65^28^.

Previously published studies using FEP for protein stability prediction all relied on the availability of experimentally resolved structures, a prerequisite that is unlikely to be true given that structures for only 15% of the human proteome have been elucidated^57^. Under circumstances where the target protein structure is unavailable, protein structure prediction approaches are needed to build a model that is suitable for FEP calculations. Homology modeling^30–33^ is one of the most widely used protein structure prediction methods and it has long been recognized that the quality of a homology model has a strong dependence on the similarity between the target and template sequence, which could in turn affect the accuracy of thermostability predictions. A previous FEP+ study to estimate protein-ligand relative binding free energies using homology models showed little to no degradation in performance when compared to predictions calculated starting from crystal structures^34^, highlighting the robustness of the FEP+ approach when an accurate structure can be utilized. The FEP+ approach has also been applied to homology models of antibody-antigen complexes, demonstrating its high accuracy in calculating the relative binding affinity of protein-protein interactions^35^.

In this study, our main goal is to understand whether models generated by homology modeling can be predictive and whether this performance is comparable to crystal structures. Following on from this, we aim to identify what effect the structural accuracy of a homology model has on FEP+ predictions and whether common errors from sequence alignment, homology model building, and FEP+ simulation might lead to inaccuracies in predictions. FEP+ calculations were performed on homology models over the same Pucci dataset as used in previous studies^20,28^, with a total of 316 mutations across 12 different protein systems. For each protein system, a varying number of homology models were built, based on the availability of homologous structures. To evaluate performance, prediction accuracy against experimental data was compared to the accuracy of calculations performed with crystal structures. Additionally, we assessed the possibility of using homology models to identify stabilizing mutants in a setting more similar to real industrial applications with the MM/GBSA-FEP+ cascade screening strategy.

## Methods

### Crystal structures dataset preparation

The Pucci *et al*. dataset^12^ was selected as the basis for this study. All data points were carefully cross-checked against the ThermoMutDB database^36^, and corrections were made to two proteins that were measured as dimers in experiments, 1AMQ and 1OH0, such that they could be simulated as monomers in this work. Two systems were excluded from the study: 1CEY was an NMR structure; whilst 1VQB had no homologous templates to be used for modeling. Mutations involving transformations between a proline or histidine residue and a charged residue were also excluded due to the intricacies involved in this type of mutation. Similarly, several mutations that required pK_a_ corrections in previous studies^20,28^ were excluded as the protonation state of these mutations would not be identifiable in homology models. A total of 316 mutations were kept covering 12 different protein systems whose structures were obtained from the Protein Data Bank^37^ and used as the baseline for evaluating the FEP+ performance (Table 1) on homology models.

**Table 1:** Performance of FEP+ predictions using homology models

| Metric | Model | Bootstrapped | Mutation data |  |  |
| --- | --- | --- | --- | --- | --- |
|  |  |  | Highest Identity<br>HM | Lowest Identity<br>HM | Highest Local<br>Identity |
| R<sup>2</sup> | xtal-FEP+ |  |  | 0.67 |  |
|  | HM-FEP+ | 0.59 ± 0.02 | 0.58 | 0.54 | 0.58 |
| RMSE<br>(kcal/mol) | xtal-FEP+ |  |  | 1.12 |  |
|  | HM-FEP+ | 1.18 ± 0.03 | 1.18 | 1.29 | 1.16 |

### Homology models dataset preparation

Homology models were constructed using the Multiple Sequence Viewer (MSV2) in BioLuminate^38^. A BLAST search for each target sequence was first performed to identify templates with identity greater than 20%. Based on protein family classification in the Pfam database^39^, only templates belonging to the same protein family as the target were considered, with at most eight templates selected per target (Table 1 and Table 5). Where there were more than eight templates available, templates were manually selected such that the global identities between the target and template had a wide range (Table 1 and Table 5). Pairwise or multiple sequence alignment was then performed for target proteins whose template sequence identities were greater than 30%. For template sequences with identities less than 30%, a combination of structural alignment and profile alignment of the templates was used to achieve better alignment results. In the structural alignment, multiple available template structures were superimposed to generate an initial sequence alignment, from which the target sequence could then be aligned. For both alignment cases, an additional manual inspection that involved protecting family-wide conserved residues and alignment of secondary structure elements based on secondary structure prediction, ensured the highest alignment quality possible. After obtaining target-template alignments, Prime^40^ was used with side chain optimization to construct all homology models using both knowledge-based and energy-based schemes^41^.

### MM/GBSA and FEP+ calculation

System preparation and calculations were performed using the 2021-2 release of the Schrödinger Suite (Schrödinger). Crystal and homology model structures were first prepared using the Protein Preparation Wizard tool in Maestro^38^. For crystal structures, all crystal waters were removed for MM/GBSA calculations, but retained for FEP+ calculations. PropKa was used to determine the protonation states of titratable residues based on the different experimental pH conditions^42^. MM/GBSA calculations were carried out using the Residue Scanning module, and FEP+ calculations were carried out using the Protein FEP+ module. The OPLS4 force field^44^ was used for all calculations.

The FEP+ methodology for single protein mutations was used as described previously^20^. Briefly, a tri-peptide composed of the mutation site residue flanked by its neighboring residues was used to model the unfolded state of the protein, with the N- and C-terminus capped by an acetyl group and an N-methyl group, respectively. All prepared systems were solvated in a 10 Å SPC water buffer. The solvated systems were subsequently equilibrated using Desmond^45,46^ with the default relaxation protocol. Perturbations were performed over 12 lambda windows for neutral mutations, 16 for proline mutations, and 24 for charge-changing mutations. Each lambda window was simulated for 10 ns; replica exchange solute tempering (REST)^47–49^ was used to enhance the sampling of mutated residues by increasing the effective temperature of the residue at the mutation site. Grand Canonical Monte Carlo^24^ was used to enhance water sampling around the mutation site.

### Local similarity analysis and bootstrapping

To evaluate how the local environment in homology models affects FEP+ prediction accuracy, local sequence identity and similarity scores were defined for each mutation site. The local 1D score was computed by counting the matches between the aligned target and template sequence in similarity for the mutation site and its six neighboring residues^50^. The local 3D score was computed by counting matching residues within 5 Å of the mutation site in the homology models. Cases where a target residue matched with a gap in the template were treated as mismatches. Matches were normalized by dividing by the total number of residues considered for each mutation site.

Since a different number of homology models were constructed for each target system, an unintended bias could be introduced when multiple data points from the same mutation site are included when reporting the overall performance of FEP+ results based on homology models. To account for this issue, a bootstrapping approach was applied, whereby for each mutation site 1000 samples of free energy predictions were drawn from the available homology models and the average performance was reported. As a result, performance metrics for FEP+ results based on homology models will be shown as bootstrapped mean ± standard deviation if applicable.

## Results

The main focus of this work was to understand how accurate thermostability predictions calculated using homology models are compared to using crystal structures. As a result, although accuracy metrics (R^2^ and RMSE) were calculated relative to the experiment, the discussion focuses on comparing these metrics for stabilities calculated with crystal structures against homology models. For convenience, the terms HM-FEP+ and xtal-FEP+ will refer to FEP+ prediction results based on homology models and crystal structures, respectively.

### Overall FEP+ performance

The performance of protein FEP+ predictions utilizing high-quality crystal structures as collected by Pucci *et al*.^12^, referred here as the “Pucci dataset”, serves as a baseline to evaluate FEP+ performance using homology models. Using the recently developed OPLS4 force field^44^, an R^2^ of 0.67 and an RMSE of 1.12 kcal/mol were achieved (Fig. 1 and Table 1). These results indicate slightly improved performance compared with the previous generation OPLS3e force field (R^2^ of 0.67 and an RMSE of 1.18 kcal/mol).

**Fig. 1:**
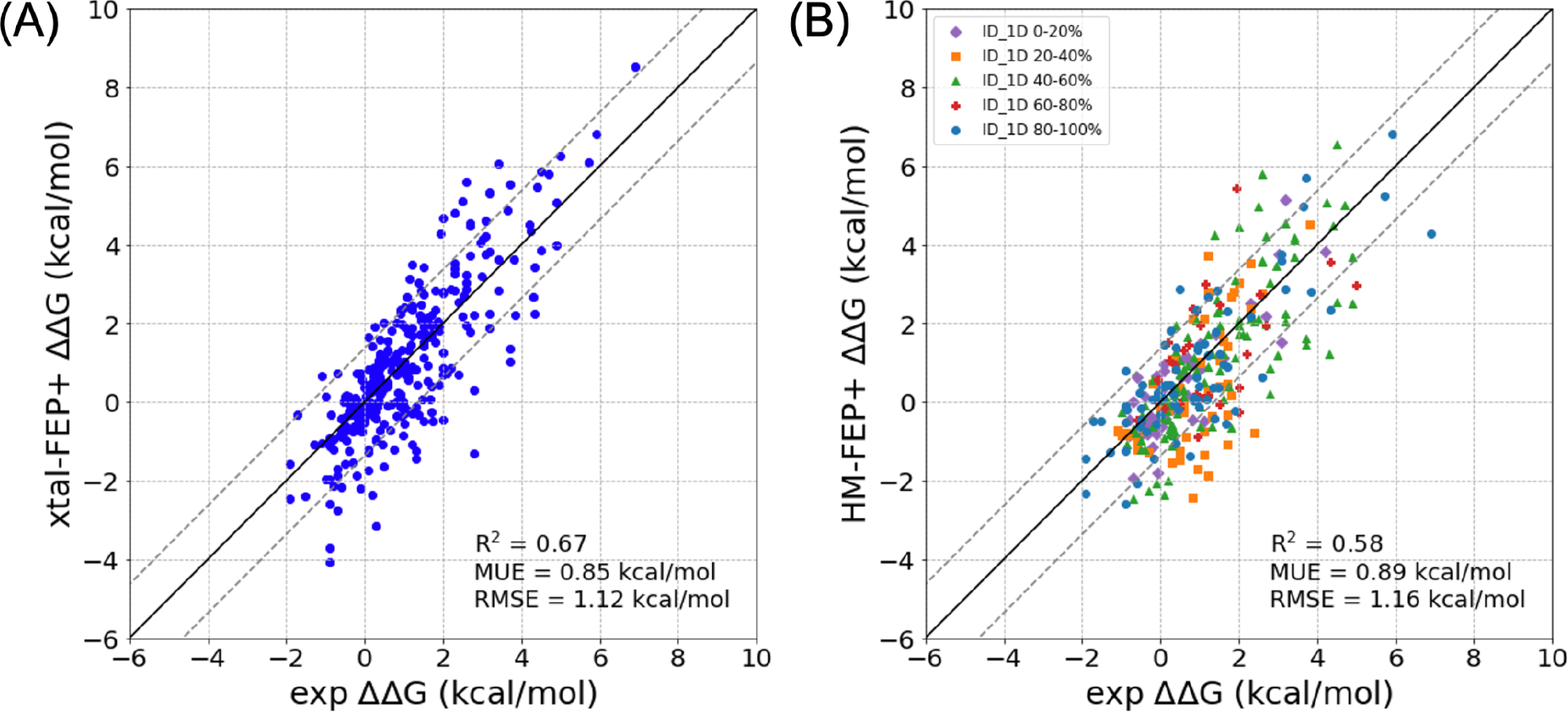
Correlation plots of FEP+ predictions against experimental data. FEP+ predictions using crystal structures (a) serve as a baseline for performance. As there were multiple templates for some systems, each mutation in the FEP+ predictions using homology models plot (b) is represented by the template that had the highest local identity score for that particular site, colored by its local 1D identity score. Dotted lines indicate an error of 1 log unit or 1.37 kcal/mol.

For the homology model dataset, a varying number of homology models were constructed for each protein system based on the availability of homologous structures (Table 2; File S1). The overall RMSEs for each system showed that certain homology models perform as well as, if not better than, the crystal structures (File S1). For the entire dataset, an R^2^ of 0.58 and an RMSE of 1.18 kcal/mol; or an R^2^ of 0.54 and an RMSE of 1.29 kcal/mol were obtained using the highest or the lowest sequence identity models for each system, respectively (Table 1 and Fig. S1). On average, an R^2^ of 0.59 and an RMSE of 1.18 kcal/mol were obtained using a bootstrapping approach (see Methods for details), indicating slightly reduced accuracy when compared with crystal structures (Table 1). The thermostability prediction errors for HM-FEP+ were found to be dependent on the accuracy of the homology model and three main factors were found to influence this accuracy: modeling schemes; unresolvable misalignments; and local quality of the template.

**Table 2:** Summary of studied systems with homology models

| Target PDB | Target protein | # of mutations | Xtal RMSE (kcal/mol) | # of HMs | Template seq. ID (%) | HM RMSE (kcal/mol) |
| --- | --- | --- | --- | --- | --- | --- |
| 1L63 | T4 Lysozyme | 112 | 1.14 | 1 | 44 | 1.11 |
| 2LZM | T4 Lysozyme | 60 | 1.06 | 1 | 44 | 1.33 |
| 1LZ1 | Human Lysozyme | 58 | 0.86 | 8 | 33 - 68 | 0.82 - 1.43 |
| 1EY0 | Staphylococcal Nuclease | 27 | 1.19 | 4 | 18 - 24 | 0.99 - 1.37 |
| 2RN2 | Ribonuclease H | 16 | 1.45 | 4 | 36 - 86 | 0.64 - 1.00 |
| 4LYZ | Hen Egg White Lysozyme | 14 | 0.91 | 5 | 35 - 84 | 0.99 - 1.17 |
| 1BNI | Barnase | 13 | 1.23 | 1 | 79 | 1.35 |
| 5PTI | Trypsin Inhibitor | 5 | 1.07 | 3 | 31 - 64 | 0.54 - 1.75 |
| 1AMQ | Aspartate Aminotransferase | 4 | 1.77 | 4 | 12 - 47 | 0.66 - 2.77 |
| 1IHB | Cyclin-Dependent Kinase 6 Inhibitor | 3 | 1.39 | 2 | 23 - 44 | 1.23 - 1.34 |
| 1OH0 | Steroid Delta-Isomerase | 2 | 1.60 | 3 | 21 - 79 | 0.97 - 1.33 |
| 1IOB | Interleukin-1 Beta | 1 | 2.68 | 5 | 19 - 78 | 1.34 - 3.11 |
| 1RN1 | Ribonuclease T1 Isozyme | 1 | 0.30 | 5 | 31 - 65 | 0.24 - 1.55 |

### Errors led by different modeling schemes in T4 lysozyme system

To highlight the effect that different modeling schemes had on the accuracy of predictions, a couple of example outliers from the bacteriophage T4 lysozyme system will be discussed. There are two target protein crystal structures for this system, 1L63 and 2LZM, which together contribute 170 mutations to the Pucci dataset. Both target proteins were modeled with the only available homologous template 7CN7, which has a global sequence identity of 44% for both targets. Two mutation sites that produced significant outliers were identified, D47 and R96/L99 (Fig. 2A and Fig. S2), whose errors could be traced back to the incorrect placement of nearby side chains in the homology models.

**Fig. 2:**
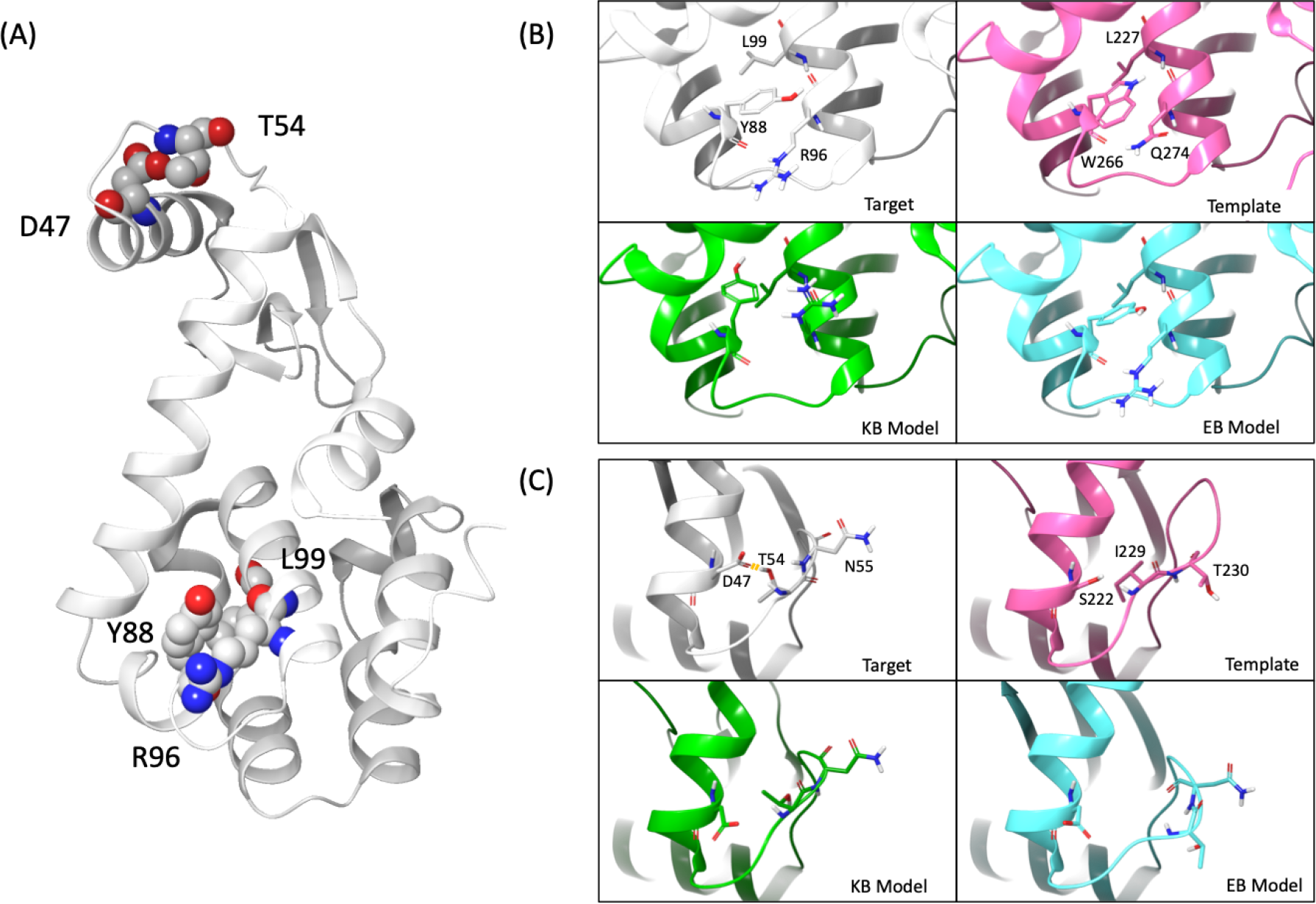
Homology models of T4 lysozyme (target sequence from 1L63), produced with either the knowledge-based (KB) or energy-based (EB) schemes using the template 7CN7. (A) Crystal structure of 1L63 highlighting the residues around two mutation sites, D47 and R96, that exhibited outliers in the predictions from homology models. (B) The KB model misplaced the Y88 side chain which led to prediction errors, whereas the EB model correctly predicted the Y88 side chain orientation. (C) The hydrogen bond between D47 and T54 was observed in the crystal structure but not in the KB or EB homology models due to the different T54 backbone conformation derived from the template.

For mutations at L99 in 1L63 and R96 in 2LZM, the side chain of Y88 was incorrectly oriented in homology models built using a knowledge-based scheme (Fig. 2B). Specifically, the side chain of Y88 occupied an apolar pocket close to L99 that was previously reported to involve ligand binding^51^ and was trapped in this local energy minimum throughout the FEP+ simulations. Furthermore, this orientation prevented interactions between Y88 and the side chain at the R96 site in both the WT and mutants. Considering all 11 mutations at R96 position, the incorrect side chain orientation of Y88 caused the FEP+ predictions using the knowledge-based homology models to produce an RMSE that compares unfavorably with the xtal-FEP+ RMSE (Table 3). By utilizing an energy-based scheme for sidechain placement, the orientation of Y88 was correctly predicted (Fig. 2B), and as a result, the RMSE for all 11 mutations improved to 1.27 kcal/mol. (Table 3). The energy-based model also improved the prediction at L99 in 1L63, in particular the L99A mutation with a thermostability prediction of 6.53 kcal/mol, which is much closer to the experimental value of 4.50 kcal/mol than the knowledge-based models (1.04 kcal/mol).

**Table 3:** Thermostability predictions (kcal/mol) for the 11 R96 mutations of the 2LZM PDB structure of T4 lysozyme with different modeling schemes

| Mutation | Exp $\Delta\Delta G$ | xtal-FEP+ $\Delta\Delta G$ | HM-FEP+ $\Delta\Delta G$<br>(knowledge-based) | HM-FEP+ $\Delta\Delta G$<br>(knowledge-based<br>w/ modified REST<br>region) | HM-FEP+ $\Delta\Delta G$<br>(energy-based) |
| --- | --- | --- | --- | --- | --- |
| R96A | 2.00 | 1.27 | 1.69 | 1.11 | 1.67 |
| R96N | 3.00 | 3.19 | 2.13 | 1.85 | 1.20 |
| R96Q | 0.30 | -0.95 | -1.60 | -0.85 | -0.91 |
| R96E | 2.50 | 5.12 | 3.73 | 3.59 | 5.26 |
| R96G | 2.60 | 2.87 | 3.46 | 3.33 | 3.56 |
| R96K | 0.00 | -0.42 | 0.22 | -0.82 | 0.22 |
| R96M | 2.70 | 1.77 | 1.91 | 1.20 | 1.91 |
| R96S | 2.60 | 3.23 | 3.60 | 3.99 | 3.39 |
| R96W | 4.50 | 3.87 | -0.95 | 2.69 | 2.85 |
| R96Y | 4.70 | 5.81 | 1.73 | 4.38 | 4.84 |
| R96V | 2.40 | 2.55 | 1.62 | 2.86 | 1.83 |
| RMSE |  | 1.05 | 2.08 | 1.11 | 1.27 |

Similarly, the D47A mutation was observed to be an outlier with an error of 4.81 kcal/mol, using the knowledge-based homology model. In the crystal structure, a hydrogen bond is formed between T54 and D47 (Fig. 2C) that cannot be formed in the D47A mutant and thus the mutant is predicted to have a thermostability of 1.98 kcal/mol in xtal-FEP+, close to the 0.95 kcal/mol observed experimentally. However, the placement of the T54 side chain in the homology model prevents this hydrogen bond from being formed in the WT (Fig. 2C), and thus the thermostability of the mutant is mispredicted to be −3.86 kcal/mol. Unlike mutations at R96/L99, the energy-based model was not able to place the side chain of T54 in the correct orientation and still resulted in an incorrect but slightly better prediction of −1.69 kcal/mol. A modified scheme where more residues were included in the REST region was able to enhance sampling to allow for T54-D47 hydrogen bond formation with the stability prediction reduced to −0.52 kcal/mol, and although still preferring the mutant, much more in line with the experimental value of 0.95 kcal/mol. In general, the energy-based scheme provided more accurate results and therefore results with this scheme will be discussed throughout the rest of the study.

### Target-template relationship and its effect on prediction accuracy in the human lysozyme system

To evaluate how the quality of a sequence alignment affects FEP+ accuracy, a thorough analysis of the human lysozyme system was performed. Eight homologous templates were identified to construct homology models including four orthologous proteins (i.e., lysozymes from other species) 2Z2F, 4LYZ, 5XUW, 3CB7 and four paralogous proteins (i.e., non-lysozyme proteins) 4YF2, 1B9O, 1ALC, 1HFX, with sequence identities ranging from 33% to 68% (Table 4 & File S1). Paralogous proteins were included as they provide further opportunities to assess whether homology models generated by proteins with different functions could still be used in FEP+ prediction, a scenario very likely to occur in practical applications.

**Table 4:** Performance of FEP+ stability predictions with different templates for the human lysozyme system

| PDB | Identity/Similarity (%) | HM C-alpha RMSD (Å) | R<sup>2</sup> | RMSE (kcal/mol) |
| --- | --- | --- | --- | --- |
| 1LZ1 | - | - | 0.67 | 0.86 |
| 2Z2F | 68/85 | 0.75 | 0.68 | 0.91 |
| 4LYZ | 60/77 | 0.73 | 0.62 | 0.96 |
| 5XUW | 52/72 | 1.45 | 0.64 | 0.83 |
| 4YF2 | 47/68 | 1.02 | 0.68 | 0.85 |
| 1B9O | 38/59 | 2.10 | 0.34 | 1.15 |
| 3CB7 | 38/53 | 1.84 | 0.70 | 0.82 |
| 1ALC | 35/58 | 1.47 | 0.69 | 0.87 |
| 1HFX | 33/54 | 1.62 | 0.31 | 1.43 |

FEP+ thermostability calculations using the 1LZ1 crystal structure produced an R^2^ of 0.67 and an RMSE of 0.86 kcal/mol when compared to the experiment. Using homology models, FEP+ predictions achieved comparable accuracy for six of the homology models with an R^2^ between 0.62 and 0.70 and an RMSE between 0.82 and 0.96 kcal/mol (Table 4). Homology models based on the two paralogs 1HFX and 1B9O crystal structures showed a reduction in accuracy with an R^2^ of 0.31 and 0.34, and an RMSE of 1.43 and 1.15 kcal/mol, respectively.

A closer examination of the two paralog templates revealed unresolvable misalignment of secondary structure features. Most outliers are located in a stretch formed by residues 102 to 115 that shows a distinct secondary structure compared to the crystal structure (Fig. 3A). In 1LZ1 (and the other homology models) residues 110 to 115 form an alpha-helix, whereas in models based on 1HFX and 1B9O, only residues 107 to 110 do. V110 showed drastically different side chain orientations in these two homology models compared to the crystal structure, which directly led to outliers including V110P/D/F/R. It is not obvious how to address such alignment issue given that there is a single residue difference between 1B9O and 1ALC (correctly modeled paralog), and residues at either end of this stretch of residues are fully conserved and well aligned across the family of proteins. To further confirm that it is the different secondary structures in the 1HFX and 1B9O that led to the poor performance, residues 102 to 115 from 1LZ1 were grafted onto 1HFX and 1B9O, and new homology models were generated. These grafted models showed significant improvements in overall accuracy, with an R^2^ of 0.62 and 0.56, and an RMSE of 0.77 and 0.94 kcal/mol, respectively.

**Fig. 3:**
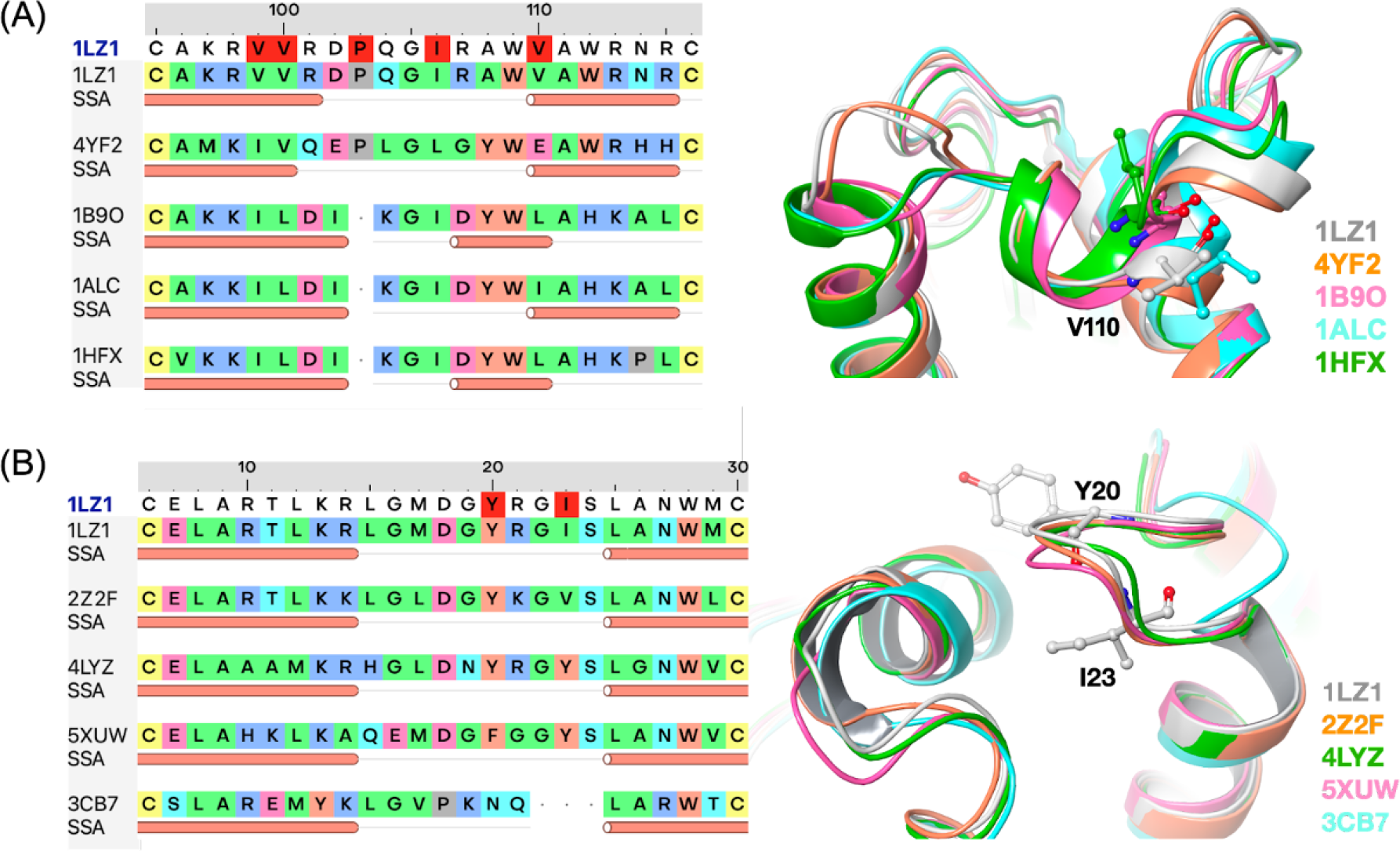
Common alignment issues leading to prediction errors in the human lysozyme system. (A) The analogous template proteins 1B90 and 1HFX displayed distinct loop conformations from the target (1LZ1), which led to errors at V110 and P103. (B) Template protein 3CB7 has a shorter loop, where two mutation sites Y20 and I23 are located, compared to the target and other template proteins. Alpha-helices are shown as red tubes below sequences. Residues in the alignment are colored by side-chain type. Target residues highlighted in red are sites of mutations.

Of the orthologs, a critical difference in the sequence alignment was identified in 3CB7 with the loop from residues 17-24 having a 3 residue deletion, leading to an incorrectly modeled loop configuration. (Fig. 3B). Two mutations, Y20F and I23A, are located in this loop, yet only I23A was an outlier in the model based on 3CB7. Using solvent-accessible surface area (SASA) as a metric for quantifying the change in the local environment, the I23A mutation has a significantly different environment in the 3CB7 model compared to the crystal structure. In the crystal structure there is a 12% reduction in the SASA for the I23A mutation but this increases to 51% for the 3CB7 model. This is in contrast to the Y20F mutation where the reduction in SASA is similar (10 vs 7%). As a result, the incorrect configuration of the loop has a limited effect on the calculated stability of the Y20F mutant, with the HM-FEP+ prediction of 0.01 kcal/mol close to both the xtal-FEP+ prediction and the experimental observation of 0.95 and 0.50 kcal/mol, respectively. The incorrect configuration has a significant impact on the I23A mutation, with the HM-FEP+ prediction of 0.15 kcal/mol being notably different from the experimental value of 2.54 kcal/mol. In an attempt to fix the I23A outlier, different alignment schemes were used to model loop 17-24, but similar predictions were still obtained ranging from 0.4 to 0.7 kcal/mol (Fig. S3) with no models built from 3CB7 able to capture the correct loop configuration observed in the crystal structure, even when the FEP+ simulation was extended to 50 ns.

### Local similarity score as a proxy for model quality validation

A number of the outliers identified throughout this study resulted from low sequence similarity around the mutation site, which is understandable given that the FEP+ method is well known to be sensitive to the local structural environment^20,25,52^. Further to this, only a small degradation in the prediction accuracy was observed when comparing the highest identity templates for each system (R^2^ of 0.58 and RMSE of 1.18 kcal/mol) to the lowest (R^2^ of 0.54 and RMSE of 1.29 kcal/mol) (Fig. S1 or Table 1), suggesting that global identity was playing a smaller role in accuracy than was expected. As a result, it is proposed that the local sequence similarity plays a more important role on the quality of modeled mutation sites and thus the FEP+ accuracy than global similarity.

To quantify this, local similarity/identity was analyzed using two metrics that take into account the residues neighboring the mutation site: a 1D or sequence similarity score; and a 3D or close (5 Å) spatial proximity score. As expected, a significant improvement in FEP+ accuracy was observed when eliminating mutations with low local scores (Table 5). For mutation sites where the local sequence identity score is above 40% or the similarity score above 60%, HM-FEP+ predictions provide comparable accuracy to xtal-FEP+ (Table 5). Surprisingly, little difference was noted between metrics calculated with the 1D and 3D scores (Table 5). This is convenient as it suggests that a homology model does not need to be generated prior to getting an indication of whether a mutation will have low or high local similarity; the 1D score will suffice. Further to this, it also means that when there are multiple templates to choose from for a single mutation site, it is possible to choose the template that has the highest 1D score for that site, rather than the template with the best overall identity. By using this selection criterion, there is an improvement in the RMSE (1.16 kcal/mol), whilst the R^2^ (0.58) is the same (Fig. 1 and Table 1), suggesting that there is a slightly better accuracy when focusing on local rather than global identity.

**Table 5:** Performance of FEP+ predictions with homology models compared to predictions with crystal structures for different local score thresholds. Performance results are shown in the format of HM-FEP+/xtal-FEP+.

| Score | $\geq 0\%$ | | $\geq 20\%$ | | $\geq 40\%$ | | $\geq 60\%$ | | $\geq 80\%$ | |
| --- | --- | --- | --- | --- | --- | --- | --- | --- | --- | --- |
| Metric | R<sup>2</sup> | RMSE (kcal/mol) | R<sup>2</sup> | RMSE (kcal/mol) | R<sup>2</sup> | RMSE (kcal/mol) | R<sup>2</sup> | RMSE (kcal/mol) | R<sup>2</sup> | RMSE (kcal/mol) |
| ID_1D | 0.59/0.67 | 1.18/1.12 | 0.57/0.65 | 1.23/1.18 | 0.63/0.67 | 1.16/1.20 | 0.62/0.72 | 1.10/1.12 | 0.67/0.77 | 1.06/1.09 |
| ID_3D | 0.59/0.67 | 1.18/1.12 | 0.60/0.68 | 1.18/1.14 | 0.64/0.67 | 1.13/1.20 | 0.61/0.67 | 1.15/1.23 | 0.57/0.66 | 1.19/1.17 |
| SIM_1D | 0.59/0.67 | 1.18/1.12 | 0.57/0.67 | 1.22/1.14 | 0.56/0.66 | 1.26/1.18 | 0.61/0.68 | 1.15/1.13 | 0.65/0.71 | 1.06/1.14 |
| SIM_3D | 0.59/0.67 | 1.18/1.12 | 0.59/0.67 | 1.19/1.13 | 0.58/0.67 | 1.23/1.17 | 0.64/0.67 | 1.13/1.18 | 0.63/0.65 | 1.10/1.22 |

To further investigate the effect of global and local identity, all mutations were categorized into four sub-groups with either high or low global and local identity (Fig S4). Based on our previous observations, mutations within a template with greater than 40% global sequence identity to the target were defined as having a high global identity, whereas those with local sequence identity above 40% were defined as having a high local identity. By analyzing the distribution of absolute errors for each group, mutations with prediction errors greater than 3 kcal/mol generally have both low global and local identity, whereas those with errors between 0-1 kcal/mol are predominantly the mutations with either high global or local identity. The overall MUE and RMSE for mutations with either high global or local identity are also smaller than those with low global and local identity (Table S1). Specifically, mutations with high local identity achieve the smallest errors among these four sub-groups. This suggests that in practice the thermostability of mutations with high local identity could be predicted with good confidence, even in cases where only a poor template could be found.

### FEP+ performance in identifying stabilizing mutations

In practice, protein stability FEP+ is predominantly utilized in protein engineering efforts to identify stabilizing mutations. To evaluate whether stabilizing mutations can be accurately predicted, the MM/GBSA-FEP+ screening cascade approach was applied as described in our previous work to the Pucci dataset^20^. Here MM/GBSA calculations are first used as a filtering step to enrich stabilizing mutation candidates for the following FEP+ predictions. True stabilizing mutations were defined as those with an experimental ΔΔG less than zero (a total of 80 out of 316 mutations), whilst stabilizing predictions had a calculated ΔΔG less than zero. Two performance metrics were calculated to assess the performance of the MM/GBSA-FEP+ cascade approach; recall (true positive rate) and precision (positive predictive value). Recall describes the percentage of true positives (TP) in the FEP+ step among all stabilizing mutations in the entire dataset, whereas precision describes the percentage of TP mutations among all stabilizing FEP+ predictions.

As there is not a one-to-one relationship between MM/GBSA energies and experimental values^53^, different cutoff values were tested ranging from 0 to 7 kcal/mol at the MM/GBSA filtering stage. As expected, an increased recall was observed with increased MM/GBSA cutoffs since more mutations were included in the FEP+ stage (Fig. S5 or Table S2). Using the MM/GBSA-FEP+ cascade approach with either crystal structures or homology models showed a slight reduction in recall (around 2-13%), corresponding to about 2 to 10 erroneous predictions among the 80 stabilizing mutations in the dataset. Surprisingly, the precision was not significantly impaired by an increased MM/GBSA cutoff, likely due to the robustness of protein FEP+ in discriminating stabilizing mutations. Comparisons to precision metrics for predictions solely made by MM/GBSA further highlights the ability of FEP+ to identify stabilizing mutations, no matter the type of initial structure (Table S2).

A common question when FEP+ is used to predict thermostability is the choice of the FEP+ cutoff for positive predictions. Recall and precision on the entire dataset were also computed for cutoffs ranging from −1 to 1 kcal/mol (Fig S6 or Table S3) with a clear tradeoff observed for different cutoffs. In general, a cutoff between −0.5 kcal/mol and 0.5 kcal/mol provides both acceptable recall and precision. Depending on experimental budgets and the number of predicted positive mutations, one may choose an FEP+ cutoff within this range for the desired recall, i.e., capturing more stabilizing mutations; and/or desired precision, i.e., making more accurate classifications.

## Discussion

### Accounting for biases present in the dataset

Our results indicate that homology models can achieve comparable accuracy to crystal structures, however, there are limitations and biases present in the dataset and results that should be discussed.

The proteins simulated in this study are all small, globular, monomeric, soluble proteins. The majority of these protein systems are enzymes; in particular the bulk of mutation data points are from the T4 and human lysozyme systems. The question of whether the accuracy of our results would remain for more complex protein systems still needs to be tested, where mutations can occur at ligand-, cofactor-, lipid- or protein-binding interfaces.

The Pucci dataset was selected for this study as it has been carefully curated to balance stabilizing mutations and destabilizing mutations. Despite this, there was still an approximately 1:3 ratio between stabilizing and destabilizing mutations across the 316 mutants. Additionally, a third of the mutations (103 mutations) have relatively small effects on protein stability (experimental ΔΔG between −0.5 and 0.5 kcal/mol). A dataset with a better balance of both stabilizing and destabilizing mutants, and with greater absolute values would allow for more confidence in the ability of the protocol to select worthwhile mutants for the experiment.

As different numbers of homology models were built for each protein system, a bias towards systems with more available templates could be introduced. In an attempt to reduce this bias and allow for a fair comparison with xtal-FEP+ predictions, a bootstrapping method was utilized such that one stability prediction was sampled for each mutation data point from the available homology models for that system. Although the bootstrapped results should indicate average performance, given multiple templates had identity below 40% and there was a decrease in performance when using low identity templates, the bootstrapped results should actually serve as a lower bound for real applications, especially when only the best homology model or models above a certain identity threshold will be considered.

### Common issues present in homology models that lead to FEP+ errors

FEP+ thermostability predictions using homology modeling demonstrates comparable accuracy to crystal structure FEP+, especially when templates with higher homology are used. From the outlier analysis described above, common issues associated with the structural accuracy of the homology models that directly lead to inaccurate FEP+ predictions were identified.

Secondary structural variability or insertions/deletions of residues, can lead to unresolved misalignment of sequences and is a major issue that can lead to low-quality homology models. Homology modeling relies on the principle that proteins with a similar sequence will have a similar structure, and yet proteins are dynamic and small changes in the sequence have been known to change the preference for a particular conformation^54^. Thus, differences in the single conformations of proteins found in crystal structures could result from the intrinsic differences in structure, but could also result from artifacts caused by specific crystallization conditions or crystal packing^55^. Gaps that cannot be resolved in an alignment will lead to secondary structures being built with non-optimal templates. If this coincides with a significant change in residue type, or a change of environment for the site of interest, such as the mutant residue being modeled as buried instead of solvent-exposed, different interactions will be formed with its neighboring residues and errors in thermostability prediction will be likely unless additional sampling can fix the misplacement. These types of issues will be hard to diagnose and will be a major limitation on the quality of FEP+ predictions associated with homology models.

Another common issue occurs when side chain conformations are erroneously modeled. There are several approaches for placing side chains in a homology model, such as knowledge-based, energy-based, and machine learning^56^. Generally, these methods will predict the same conformation of a side chain but there are times when misplaced side chains can lead to a local structure rearrangement and as a result, incorrect thermostability predictions. Currently, there is no good way to understand *a priori* which method is best for a particular protein, let alone a particular region of a protein.

Most of these issues occur when a poor homologous template is used for the particular site of interest, and as a result, the target mutation site is then modeled with low confidence. Despite this, by understanding the issues mentioned above it is possible to get an indication as to how much confidence should be placed in a thermostability prediction for a specific site. In particular, one should ensure that they have the highest quality homologous protein as a template; obtain as much familial sequence information as possible to obtain the best alignment; understand the likely secondary structure around the mutation site, and whether that site occurs at a region of high structural/conformational variability; calculate the local conservation score for each site of interest; use a virtual screening cascade with appropriate cutoffs for the number of FEP+ calculations you want to run, and the number of mutants you would eventually like to produce/synthesize.

## Conclusion

Given a high homology template for the regions of interest in a protein, the absence of a crystal structure is no longer a major roadblock when predicting thermostabilities. The results presented here agree with the conclusions from recent studies where thermostabilities were calculated for models generated by AlphaFold and homology modeling^56,58,59^. To get results consistent with the experiment, the authors suggest using templates with at least 40% sequence identity. In our study, we were able to show that this 40% identity need not be at the global but the local level. By using templates where the global sequence identity or local 1D sequence identity is greater than 40%, FEP+ results on homology models have an overall RMSE of less than 1.1 kcal/mol, comparable to results using crystal structures. When utilized as a part of an *in silico* screening cascade with MM/GBSA, this allows large numbers of mutants to be screened quickly and accurately.

The workflow discussed here should be utilized first to understand whether a particular template will allow for a homology model to be built that will allow for accurate thermostability predictions for a set of sites of interest, and then for performing the actual calculations. A number of further validations and enhancements are envisioned to increase the accuracy and expand the domain of applicability of this workflow. These include understanding whether running predictions on an ensemble of conformations of a homology model, as produced by molecular dynamics or by generating multiple models, could lead to more accurate predictions; being able to predict whether disulfide bridges could be formed and their effect on stability; confirming the ability of this workflow to deal with multiple mutants (double, triple, etc.) especially non-additive multiple mutations. Going forward it is hoped that this workflow could eventually be adapted to also search for potential sites of mutation that have a high likelihood of leading to increases in thermostability.

## Acknowledgments

We thank Agustina Rodriguez-Granillo, Jianxin Duan, and Antonija Kuzmanic for their useful discussions.

