## Supporting Information for "Target-template relationships in protein structure prediction and their effect on the accuracy of thermostability calculations"

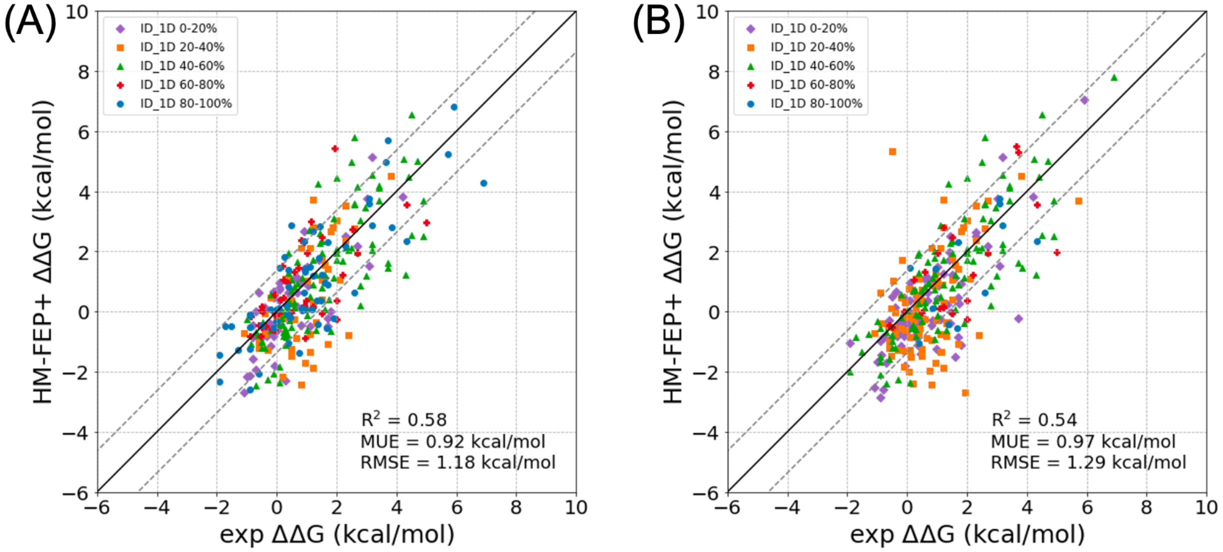


Fig. S1: Correlation plots between FEP+ predictions against experimental data using the best (a) or the worst (b) HMs for each system, colored by their local 1D identity score.

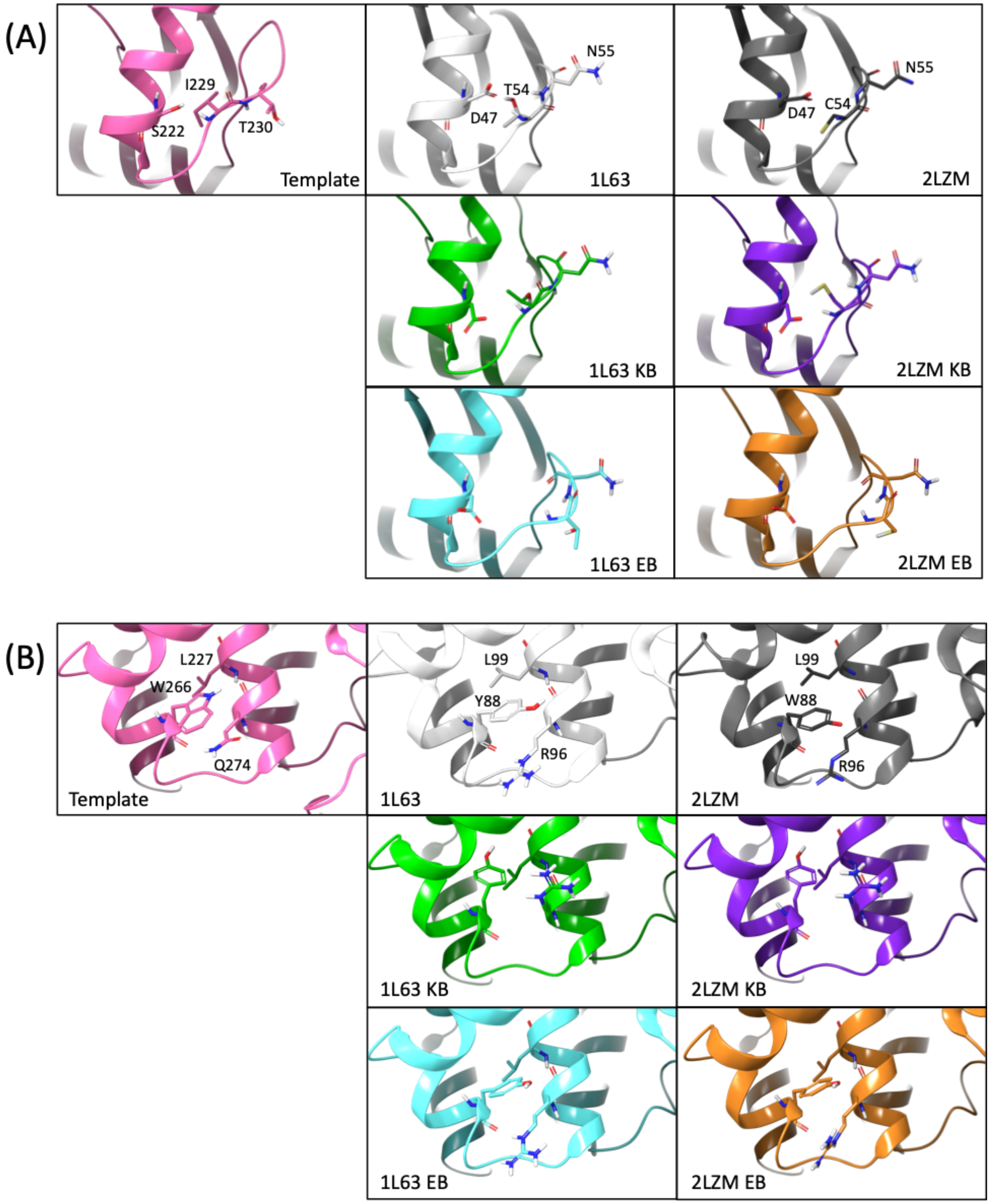


Fig. S2: Structures of the template (7CN7), the 1L63 and 2LZM targets, and the knowledge-based (KB) and energy-based (EB) models around the (A) D47 and (B) R96 mutation site, highlighting the similarities between the models built for different targets

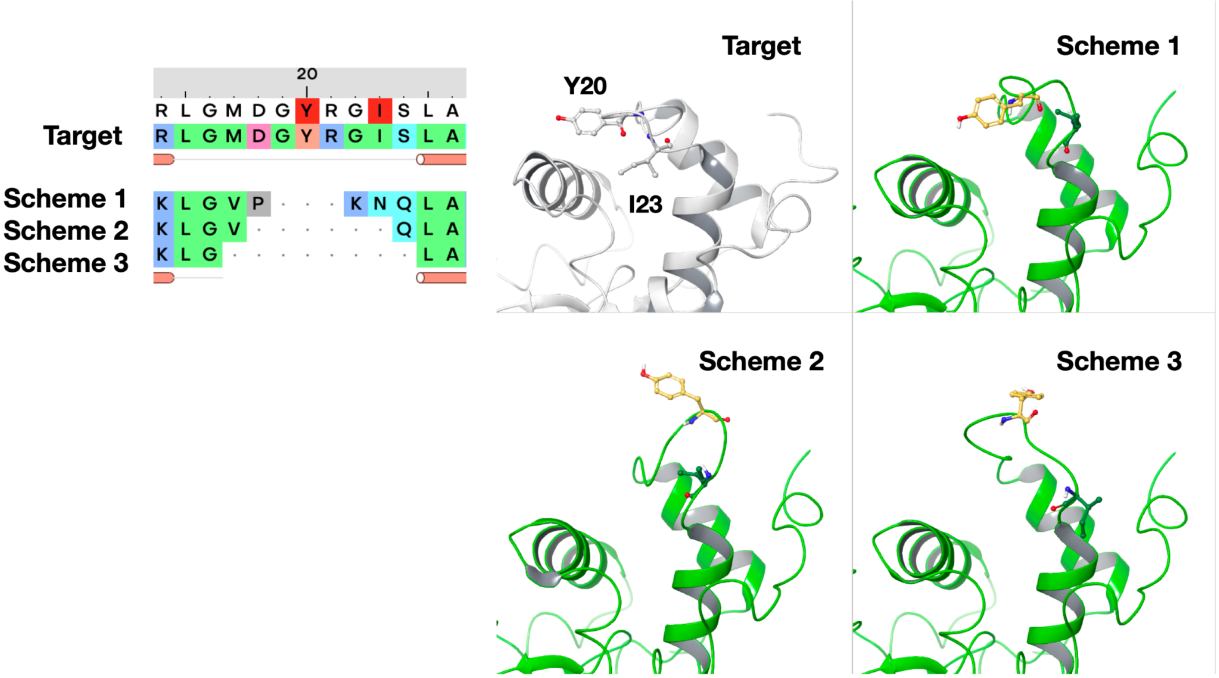


Fig. S3: Different alignments tested for loop 15-24 in homology modeling of human lysozyme using 3CB7 as the template. For the outlier mutation I23A (experimental ΔΔG = 2.5 kcal/mol), both schemes 1 and 2 predicted a thermostability change of 0.7 kcal/mol and scheme 3 predicted a change of 0.4 kcal/mol.

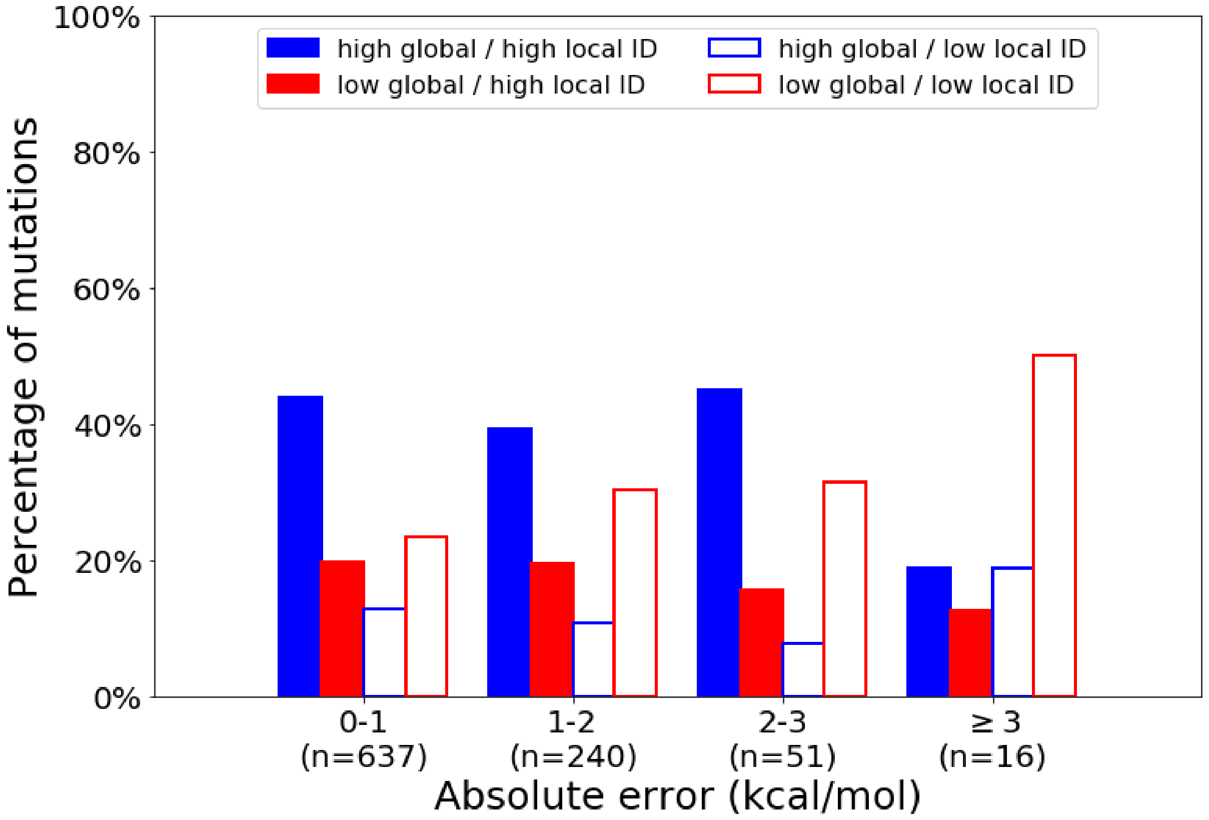


Fig S4: Distribution of prediction errors for mutations with high or low global/local identity. Normalization is performed within each error category. The cutoffs for both global and local identities are 40%.

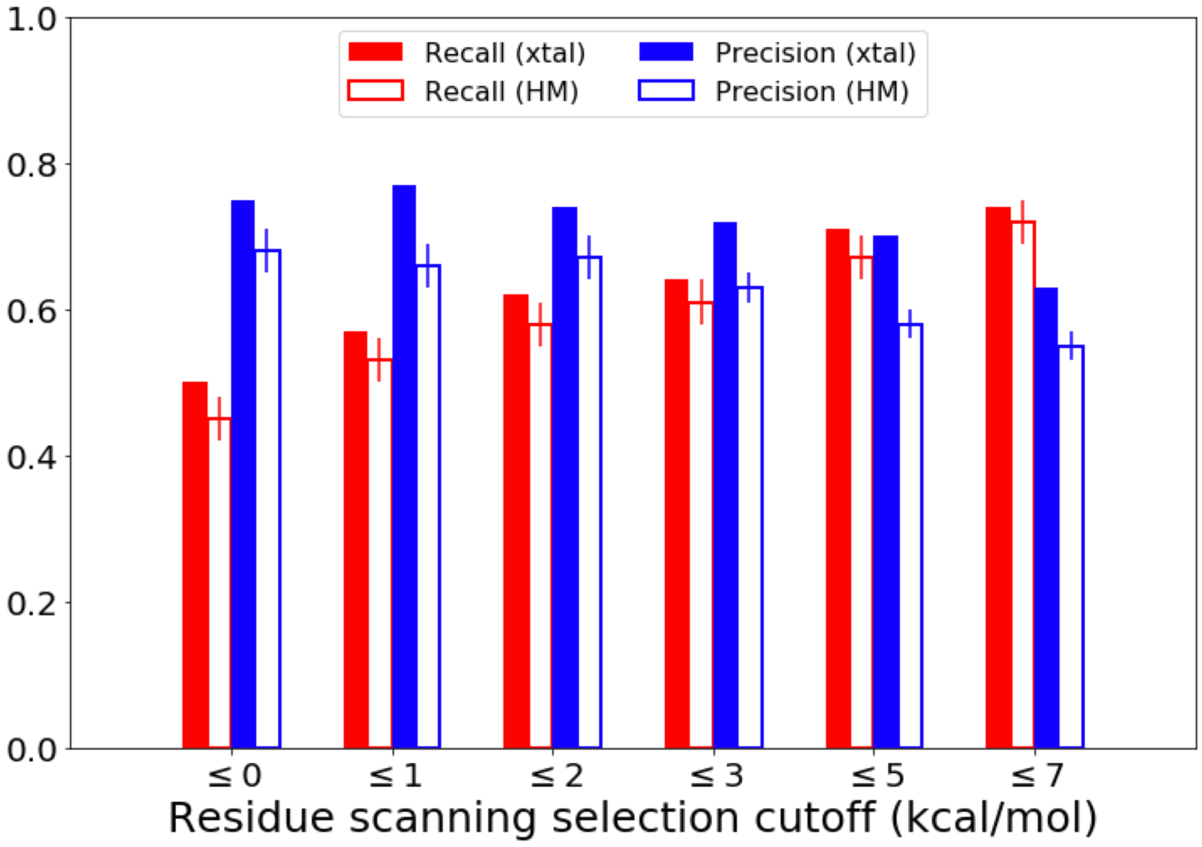


Fig. S5: Classification performance of MM/GBSA-FEP+ cascade screening using either crystal structures (xtal) or homology models (HM) with different MM/GBSA cutoffs for stabilizing mutations.

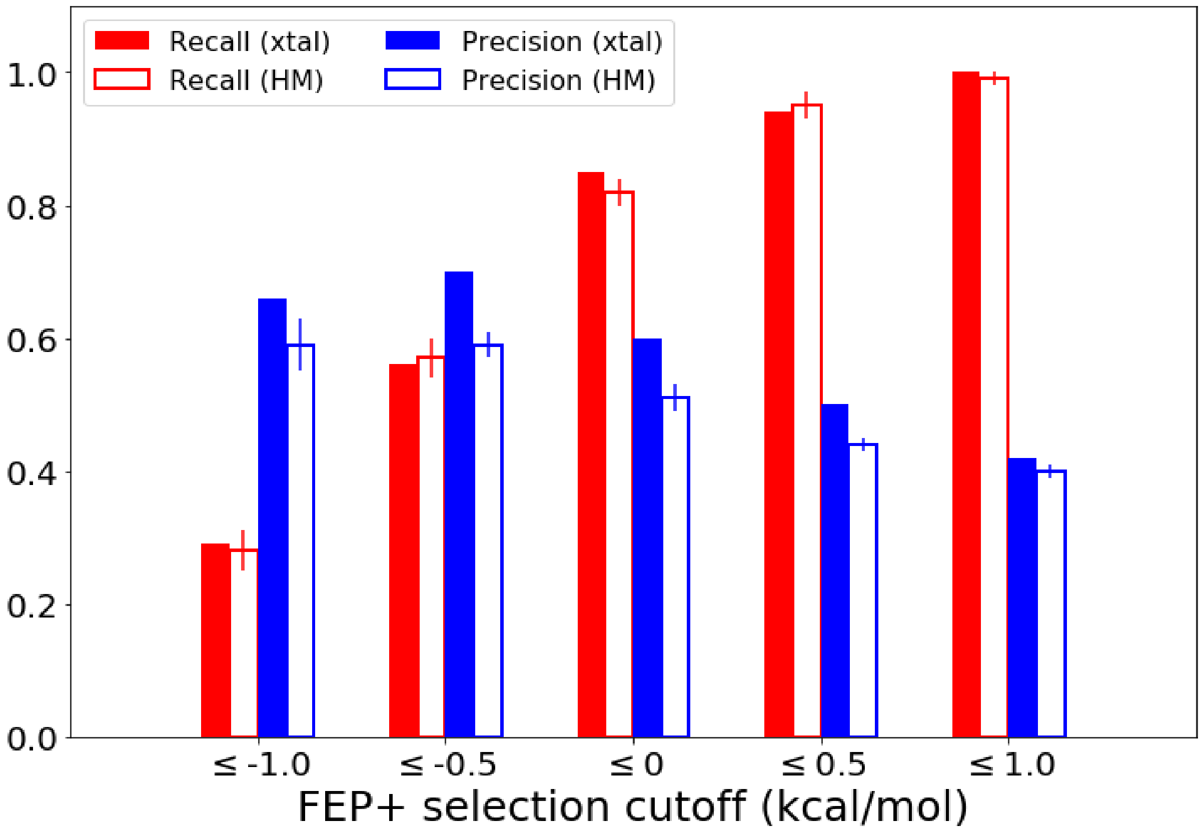


Fig. S6: Classification performance using either crystal structures (xtal) or homology models (HM) with different FEP+ cutoffs for stabilizing mutations over the entire dataset.

Table S1: Prediction errors for mutations with high or low global/local identity. The cutoffs for both global and local identities are 40%.

| | | Low Local | High Local |
|---|---|---|---|
| Low Global | MUE (kcal/mol)<br>RMSE (kcal/mol) | 0.99<br>1.33<br>(n = 246) | 0.80<br>1.03<br>(n = 183) |
| High Global | MUE (kcal/mol)<br>RMSE (kcal/mol) | 0.83<br>1.09<br>(n = 115) | 0.82<br>1.04<br>(n = 400) |

Table S2: Classification performance of the MM/GBSA-FEP+ cascade screening using either crystal structures or homology models. Standard deviations of the bootstrapped recall and precision values for homology modeled are all below 0.03.

| | | Selected MM/GBSA cutoff/filtering threshold (kcal/mol) | | | | |
|---|---|---|---|---|---|---|
| Metric | Model | ≤ 0 | ≤ 1 | ≤ 3 | ≤ 5 | ≤ 7 |
| Recall | xtal-MM/GBSA | 0.55 | 0.66 | 0.75 | 0.82 | 0.85 |
| | xtal-FEP+ | 0.50 | 0.57 | 0.64 | 0.71 | 0.74 |
| | HM-MM/GBSA | 0.54 | 0.62 | 0.72 | 0.79 | 0.85 |
| | HM-FEP+ | 0.45 | 0.53 | 0.61 | 0.67 | 0.72 |
| Precision | xtal-MM/GBSA | 0.53 | 0.55 | 0.48 | 0.42 | 0.38 |
| | xtal-FEP+ | 0.75 | 0.77 | 0.72 | 0.70 | 0.63 |
| | HM-MM/GBSA | 0.51 | 0.49 | 0.46 | 0.40 | 0.36 |
| | HM-FEP+ | 0.68 | 0.66 | 0.63 | 0.58 | 0.55 |
| # of FEP+ simulations | xtal | 83 | 96 | 125 | 158 | 179 |
| | HM | 85 ± 3 | 102 ± 4 | 126 ± 4 | 159 ± 3 | 187 ± 3 |

Table S3: Classification performance using either crystal structures (xtal) or homology models (HM) with different FEP+ cutoffs for stabilizing mutations.

| Metric | Model | Selected FEP+ cutoff/filtering threshold (kca/mol) | | | | |
|---|---|---|---|---|---|---|
| | | ≤ -1 | ≤ -0.5 | ≤ 0 | ≤ 0.5 | ≤ 1 |
| Recall | xtal-FEP+ | 0.29 | 0.56 | 0.85 | 0.94 | 1.00 |
| | HM-FEP+ | 0.28 ± 0.03 | 0.57 ± 0.03 | 0.82 ± 0.02 | 0.95 ± 0.02 | 0.99 ± 0.01 |
| Precision | xtal-FEP+ | 0.66 | 0.70 | 0.60 | 0.50 | 0.42 |
| | HM-FEP+ | 0.59 ± 0.04 | 0.59 ± 0.02 | 0.51 ± 0.01 | 0.44 ± 0.01 | 0.40 ± 0.01 |
